# Estrogen is Coupled to the Calcium Release Machinery in Human Leiomyoma Cells

**DOI:** 10.64898/2026.09.22.753480

**Authors:** Ariana Machado, Jasmine Kaur, Veronica Kan, Ying Fan, Mostafa Borahay, Darren Boehning

**Author notes:** These authors contributed equally (Ariana Machado, Jasmine Kaur).

## Abstract

Uterine leiomyomas, commonly known as fibroids, are benign tumors of the female reproductive tract. They are the most common tumor of the female reproductive system and the leading cause of hysterectomy in the United States. Progesterone, estrogen, and their receptors are important drivers for the development and growth of leiomyomas. How estrogen drives leiomyoma growth remains unclear. We have previously shown that simvastatin can inhibit leiomyoma growth by modulating intracellular calcium. This suggests that calcium signaling is important for leiomyoma growth. We therefore hypothesized that estrogen may drive leiomyoma growth by stimulating calcium release through the membrane estrogen receptor GPR30 (also known as GPER). We found that estrogen dose-dependently induced calcium release in leiomyoma cells. The GPR30-specific agonist G-1 also induced calcium release, indicating that GPR30 mediates estrogen-mediated calcium release. The GPR30 antagonist G-15 blocked estrogen-induced calcium release and significantly reduced leiomyoma proliferation. Our results demonstrate that GPR30 is a major driver of estrogen-dependent growth in leiomyoma cells by modulating intracellular calcium. These findings may have high therapeutic relevance to leiomyomas and other estrogen-dependent tumors.

## INTRODUCTION

In women of reproductive age, uterine leiomyomas, or fibroids, are a common benign tumor that occurs in the uterine wall (1). These hormonally driven tumors often cause chronic pain, significant menstrual bleeding, anemia, and infertility. Leiomyomas are the leading cause of hysterectomy in the United States, underscoring both their prevalence and the limited success of current medical therapies (2). Despite the high disease burden, the molecular mechanisms that drive leiomyoma growth remain incompletely understood. Leiomyomas are hormonally responsive, with both progesterone and estrogen acting as key regulators of tumor growth and extracellular matrix expansion (1).

Estrogen signaling has been a longstanding target for therapies geared toward leiomyoma growth management. Classical estrogen actions occur through nuclear estrogen receptors, which function as ligand-activated transcription factors to regulate gene expression (1, 3). However, estrogen also signals through non-genomic, rapid response pathways initiated by membrane receptors. These faster responses are mediated by membrane-associated estrogen receptors, including GPR30/GPER, which initiate non-canonical estrogen signaling (4). Through G protein coupling, GPR30 can activate second-messenger pathways such as inositol 1,4,5-trisphosphate (IP_3_) signaling, which can alter intracellular calcium within seconds to minutes. Although GPR30 has been associated with several estrogen-responsive tissues and cancers, its role in leiomyomas has not been fully investigated (5-9). This represents an important knowledge gap in the field, given the therapeutic limitations of systemic estrogen suppression. Aromatase inhibitors and other estrogen-decreasing agents can reduce leiomyoma growth and volume, but their use is associated with side effects such as bone loss, vasomotor symptoms, and cardiovascular risk (1). Identifying membrane-initiated estrogen mechanisms may consequently reveal alternative, more targeted approaches and therapies for leiomyoma management with fewer side effects.

Calcium signaling has recently emerged as a critical regulator of leiomyoma cell survival (10-12). Prior work from our group demonstrated that simvastatin potently inhibits leiomyoma proliferation and induces apoptosis through a calcium-dependent mechanism (11). Specifically, simvastatin treatment decreased ERK phosphorylation, altered cell cycle progression, and triggered robust apoptotic signaling that required calcium release. This response was accompanied by elevated cytosolic and mitochondrial calcium, mitochondrial depolarization, and increased caspase-3 activation. Importantly, chelation of intracellular calcium with BAPTA completely abolished simvastatin-induced apoptosis, and pharmacologic blockade of voltage-gated calcium channels prevented cell death, indicating that calcium influx through L-type channels is essential for this process (11). These findings reveal that the apoptotic machinery in leiomyoma cells is tightly coupled to intracellular calcium handling. Based on this work, we hypothesized that estrogen may also drive leiomyoma growth by directly engaging the calcium-release machinery via upstream membrane-initiated signaling.

Here, we tested the hypothesis that estrogen promotes leiomyoma growth by activating calcium release through the membrane estrogen receptor GPR30. We found that human leiomyoma (HuLM) cells exhibit a strong, dose-dependent calcium release in response to estrogen, and that this response can be replicated by selective activation of GPR30 with the selective agonist G-1. The selective GPR30 antagonist G-15 potently blocks estrogen-induced calcium release and leiomyoma cell proliferation. These results indicate that GPR30 warrants further investigation as a therapeutic target to limit leiomyoma growth.

## MATERIALS AND METHODS

### Cell culture

The immortalized human leiomyoma cell line (HuLM) was a gift from Dr. A. Salama (Baylor College of Medicine, Houston, TX). In this cell line, human leiomyoma cells obtained from a patient after surgery were immortalized using a retroviral vector carrying human telomerase reverse transcriptase (13). HuLM cells were maintained in smooth muscle cell growth medium (Lonza, Walkersville, MD), containing smooth muscle basal medium, 5% fetal bovine serum, 0.1% insulin, 0.2% recombinant human fibroblast growth factor B, 0.1% of a mixture of gentamicin sulfate and amphotericin B, and 0.1% human epidermal growth factor in 5% CO2 at 37 °C.

### Reagents and Chemicals

17β-estradiol (E2) was purchased from Sigma-Aldrich, and a 10 mM stock solution was prepared in 100% ethanol. The selective GPR30 agonist G-1 was purchased from VWR, and the antagonist G-15 was obtained from Cayman Chemical. Both G-1 and G-15 were dissolved in DMSO as stock solutions. Anti-GPR30 was obtained from Thermo Fisher Scientific. All other chemicals and reagents used in the experiments were purchased from Thermo Fisher Scientific and Sigma-Aldrich.

### Immunofluorescence Imaging of GPR30

Super-resolution imaging: Human leiomyoma cells were plated on coverslips in a 6 well plate. After 24h, the cells were fixed using 4% ice-cold paraformaldehyde at RT with rotation for 20 minutes. The cells were then quenched with 30 mM glycine/PBS solution for 5 minutes at RT with rotation. Next, the cells were washed 3 times with PBS. Cells were permeabilized with 0.25% Triton X-100 and 1% BSA in PBS for 10 minutes at RT with rotation. The primary GPR30 Recombinant Rabbit Monoclonal Antibody (20H15L21) was purchased from Invitrogen and used in 1:100 dilution. After Cells were incubated with primary antibody for 1h in RT, they were blocked with 2% BSA in PBS for 1 hour. Cells were washed 3 times with PBS for 5 minutes each and then incubated with anti-rabbit Star Red secondary antibody (1:500) obtained from Abberior Inc in PBS with 0.3% BSA for 1h. Then, cells were washed 3 times with PBS. Finally, cells were mounted on glass slides sealed with nail polish. STED super-resolution imaging was performed using a STEDYCON microscope (Abberior) using 100X objective.

### Fura-2 Calcium Imaging

HuLM cells were serum-starved for 24 hours prior to calcium imaging. Starved cells were loaded with 1 μM Fura-2 AM dye for 30 minutes at room temperature in imaging solution as described (11). Following dye loading, cells were washed and maintained in imaging solution without dye for an additional 20 minutes before imaging. Fluorescence imaging was performed using a Nikon widefield microscope equipped with a 40X objective and Elements acquisition software. Images were acquired every 2 seconds with excitation at 340 nm and 380 nm and emission at 510 nm. Baseline recordings were captured for 1 minute before the addition of 17β-estradiol (E2), G-1, or vehicle, after which imaging continued for a total duration of 15 minutes. Where indicated, cells were pre-incubated with the GPR30 antagonist G-15 prior to estrogen stimulation. Calcium imaging data were analyzed using Microsoft Excel and GraphPad Prism software. Regions of interest (ROIs) were manually selected, and fluorescence intensity traces were generated for each ROI. Calcium release events were identified as transient peaks in fluorescence ratio that returned to baseline. For each event, the following parameters were recorded: time of first release, time of peak release, release amplitude (peak fluorescence ratio minus baseline ratio immediately prior to the peak), and number of release events per cell. The number of total responding cells was also quantified. Cells exhibiting spontaneous calcium release prior to agonist addition were excluded from analysis. Release events with amplitudes below 0.01 ratio units were also excluded. Summary statistics, including responder percentages and release kinetics, were calculated, and data visualization was performed using GraphPad Prism. Dose-response data in Figure 3 were statistically analyzed using a one-way ANOVA. Percent responders in Figure 4 were calculated using an unpaired two-tailed t-test. Comparison groups were 1 μM G-1 versus 10 μM G-1; E2 1 μM versus E2 1 μM + G15; E2 1 μM versus E2 1 μM in zero-calcium solution.

### MTT Assay

HuLM cells were seeded at 5,000–7,500 cells/well in 96-well plates in 100 µL of growth medium. After 24 hours, the medium was replaced with serum-free starvation medium. Following 24 hours of serum starvation, cells were treated with the indicated concentrations of the compound (0, 0.1, 1, 2, 5, and 10 µM) diluted in the starvation medium (100 µL/well). After 48 hours of treatment, 10% (v/v) MTT reagent (5 mg/mL in starvation medium) was added, and cells were incubated at 37°C for 1 hour. The media was then removed, and formazan crystals were solubilized in 100 µL of DMSO. Absorbance was measured at 570 nm with background correction at 690 nm. Statistical significance was determined by comparing vehicle to each individual concentration using an unpaired two-tailed t-test.

## RESULTS

### HuLM cells express the membrane estrogen receptor GPR30

To investigate the role of membrane estrogen receptor signaling in leiomyomas, we first examined the expression and subcellular localization of GPR30 in immortalized human uterine leiomyoma (HuLM) cells. Immunofluorescence staining revealed robust GPR30 expression throughout HuLM cells (**Figure 1**). Superresolution microscopy analysis demonstrated that GPR30 exhibited a predominantly intracellular distribution pattern, with notable accumulation in the perinuclear region and cytoplasm. This intracellular localization is consistent with recent reports indicating that GPR30 can reside on intracellular membranes, including the endoplasmic reticulum and Golgi apparatus, in addition to the plasma membrane (4, 14). Our results are also consistent with other studies showing overexpression of GPR30 in leiomyoma tissue when compared to normal myometrial tissue (9). The strong expression of GPR30 in HuLM cells suggested that this receptor may play a functional role in mediating estrogen signaling in leiomyoma tumors.

**Figure 1.**
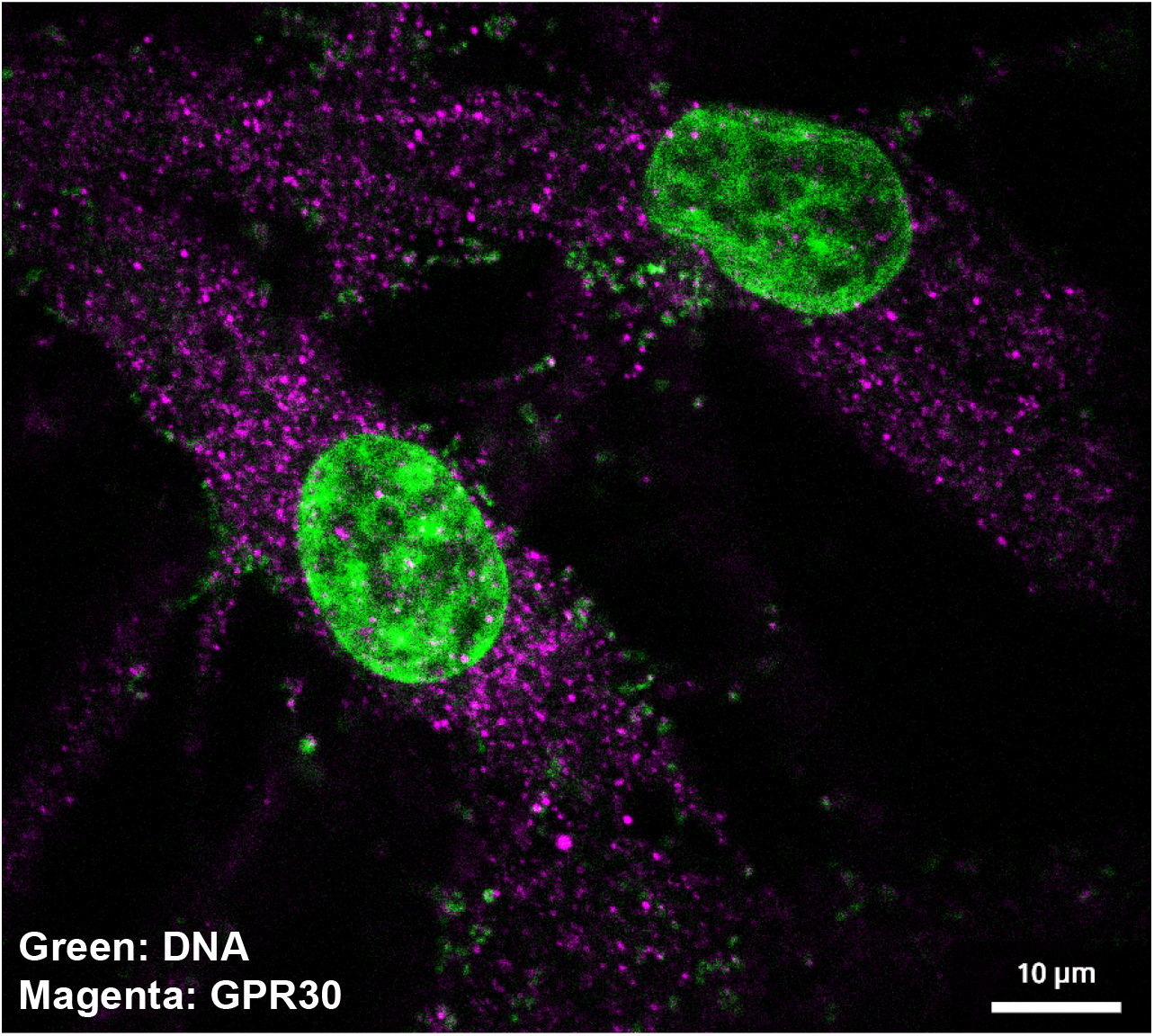
Immunofluorescence staining of GPR30 in human leiomyoma cells. Human leiomyoma (HuLM) cells were stained with anti-GPR30 (magenta/purple) and BioTracker 488 Green nuclear dye (green) and imaged using STED superresolution microscopy. GPR30 is abundantly expressed throughout HuLM cells, displaying a predominant cytoplasmic distribution pattern.

### Estrogen Stimulates Dose-Dependent Calcium Release in HuLM Cells

Given the established importance of calcium signaling in leiomyoma cell survival (10-12) and our observation of GPR30 expression, we hypothesized that estrogen regulates intracellular calcium dynamics through GPR30-mediated signaling. To test this hypothesis, we performed live-cell Fura-2 calcium imaging. 17β-estradiol (E2) treatment induced robust calcium release in HuLM cells in a dose-dependent manner (**Figures 2,3**). Representative Fura-2 ratio images captured at 1, 3, 5, 7, and 9 minutes following E2 addition demonstrated clear calcium elevation in responding cells (**Figure 2A; Supplementary Movie 1**). Single-cell calcium traces revealed that E2 triggered rapid calcium release events characterized by a sharp increase in intracellular calcium concentration followed by a return toward baseline levels (**Figure 2B**). The calcium response was observed within seconds to minutes following E2 application, consistent with a non-genomic, membrane-initiated signaling mechanism. Quantitative analysis of calcium release revealed a clear dose-response relationship (**Figure 3**). At low E2 concentrations (10 nM), approximately 20-30% of HuLM cells exhibited calcium release. As the E2 concentration increased to 100 nM and 1 μM, the percentage of responding cells progressively increased to approximately 40-50% and 60-70%, respectively. At the highest concentration tested (10 μM E2), nearly 80-90% of cells demonstrated calcium release. These findings establish that estrogen potentially stimulates calcium mobilization in leiomyoma cells and that this response exhibits a dose-dependent pattern consistent with receptor-mediated signaling.

**Figure 2.**
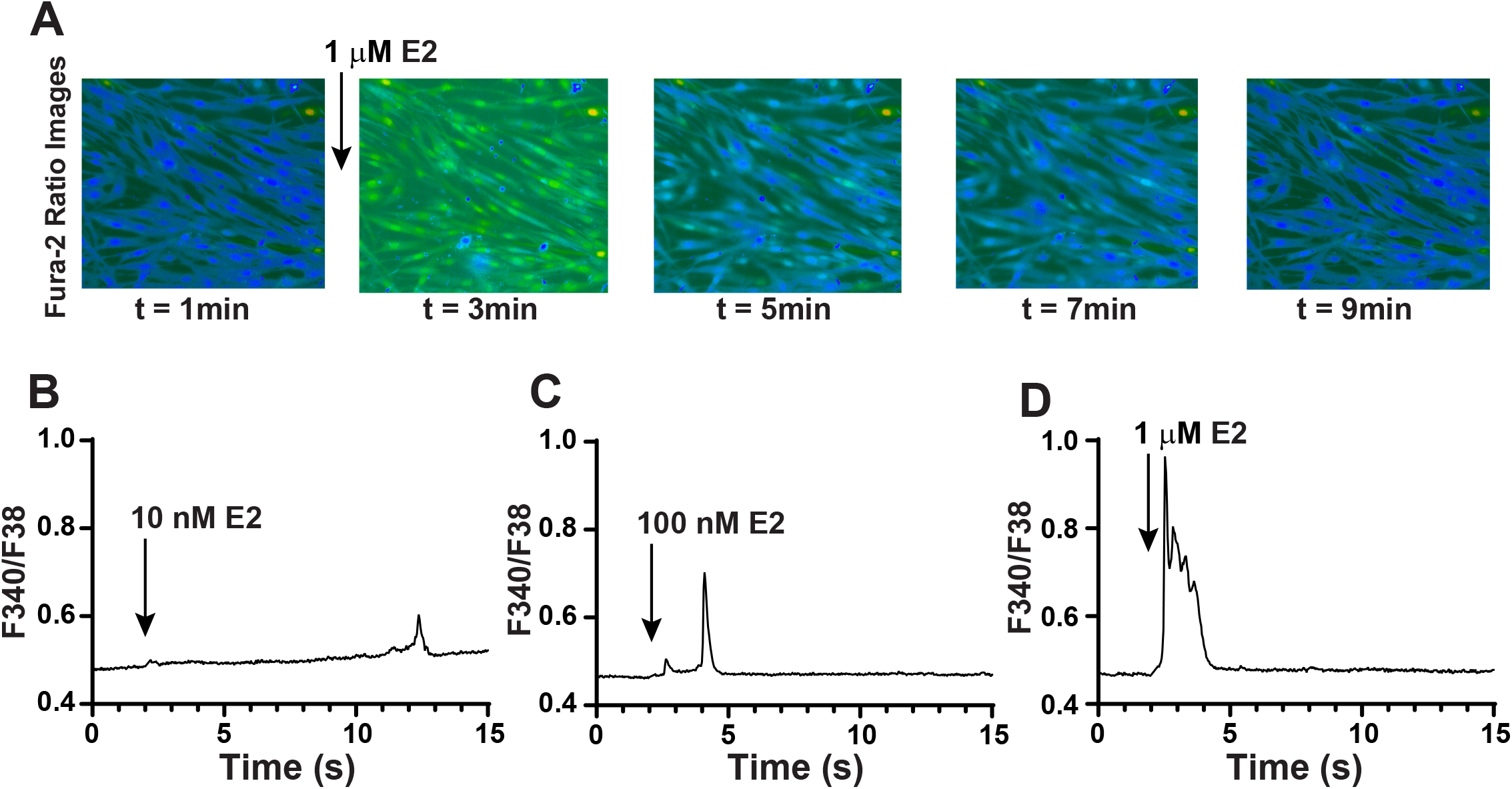
17β-estradiol (E2) induces calcium release in HuLM leiomyoma cells. **(A)** Fura-2 ratio images of living HuLM cells treated with 1 µM E2 at 2 minutes, showing real-time changes in intracellular calcium over time (t = 1, 3, 5, 7, and 9 minutes). Color shifts from blue (baseline) to green/yellow indicate elevated cytosolic calcium concentration. **(B–D)** Representative Fura-2 time-series single-cell traces demonstrating rapid cytosolic calcium release in HuLM cells treated with indicated doses of E2 (B: 10 nM, C: 100 nM, D: 1 µM) added at 2 minutes (arrows).

**Figure 3.**
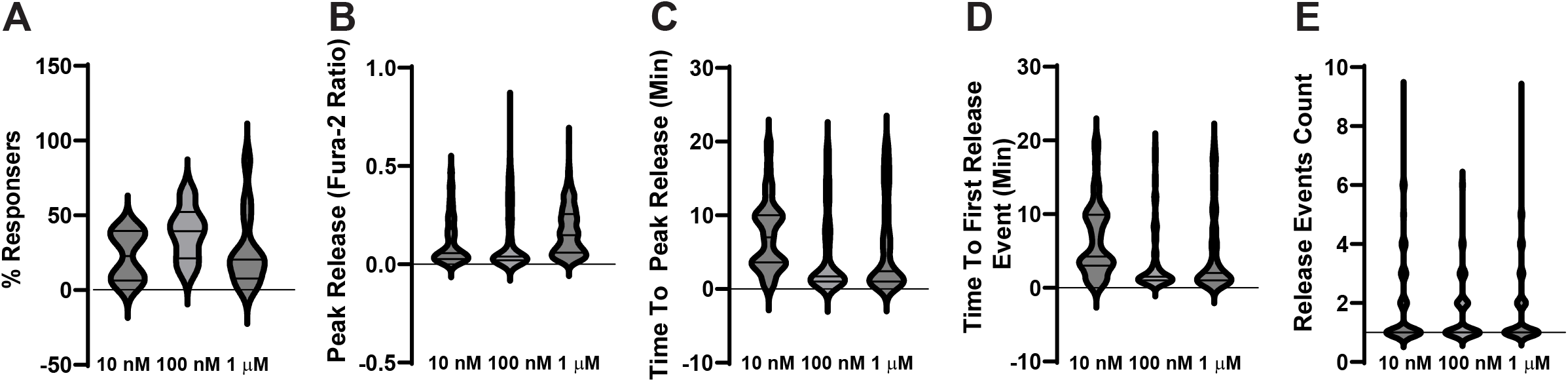
Calcium dynamics in HuLM cells at varying doses of 17β-estradiol (E2). **(A)** Percentage of Responders: Proportion of analyzed HuLM cells (% Responders) exhibiting calcium release following application of 10 nM, 100 nM, and 1 µM E2. **(B)** Peak release amplitude (ratio units): Maximum calcium response expressed as the peak Fura-2 fluorescence ratio at each indicated E2 concentration. **(C)** Time to Peak Release Amplitude: Time elapsed (minutes) from E2 treatment addition to maximum cytosolic calcium concentration. **(D)** Time to first calcium release event (minutes): Onset latency (minutes) from E2 application to the initial intracellular calcium spike. **(E)** Release Events Count: Total number of distinct intracellular calcium release spikes observed during the 30-minute recording period. All panel graphs were statistically significant as determined by ANOVA (*P<0*.*01*). The data represent at least 5 independent experiments comprising hundreds of single-cell traces.

### GPR30 Mediates Estrogen-Induced Calcium Release in HuLM Cells

To determine whether GPR30 specifically mediates the estrogen-induced calcium response, we employed G-1, a selective GPR30 agonist that does not activate classical nuclear estrogen receptors ERα or ERβ (15). Treatment of HuLM cells with G-1 recapitulated the calcium release response observed with E2 (Figure 4B). G-1 induced dose-dependent calcium release, with increasing concentrations producing progressively greater percentages of responding cells. At 100 nM G-1, approximately 40-50% of cells exhibited calcium release, while 1uM G-1 elicited responses in approximately 60-70% of cells. At 10 μM G-1, the response rate approached 80-90%, closely mirroring the maximal response observed with E2. The striking similarity between E2- and G-1-induced calcium release patterns strongly suggest that GPR30 is the primary receptor mediating estrogen-stimulated calcium mobilization in leiomyoma cells. The rapid kinetics of the response, occurring within seconds to minutes of agonist addition, further support a non-genomic, membrane-initiated signaling mechanism rather than transcriptional regulation through nuclear estrogen receptors.

**Figure 4.**
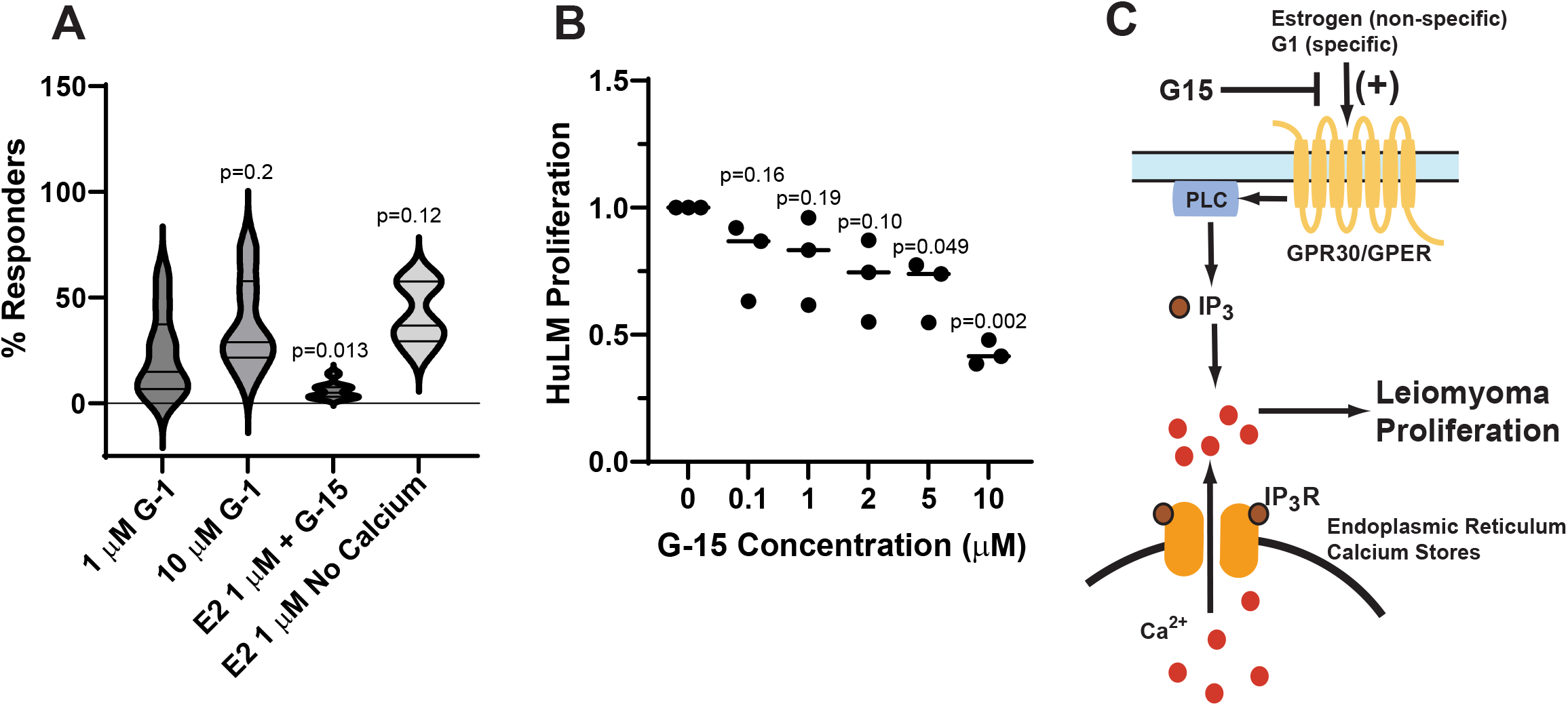
17β-estradiol (E2)-induced calcium release and cell proliferation are mediated by GPR30. **(A)** Percentage of responding HuLM cells (% Responders) exhibiting calcium release following treatment with selective GPR30 agonist G1 (1 µM and 10 µM), 1 µM E2 combined with GPR30 antagonist G-15, or 1 µM E2 in calcium-free imaging solution. There was no statistical difference between 1 μM and 10 μM G-1 (p = 0.2). G-15 preincubation significantly inhibited 1 μM E2 responses (comparison group is Figure 3A, 1 μM; p=0.013). The % responders in calcium-free imaging solution did not differ significantly from responses in calcium-replete media (comparison group is Figure 3A, 1 μM; p=0.12). P-values were calculated relative to comparison groups using an unpaired two-tailed t-test. The data reflect at least 5 independent experiments from hundreds of single-cell traces. **(B)** Relative HuLM cell proliferation assessed by MTT assay after 48 hours of treatment with indicated concentrations of G-15 (0 to 10 µM). Horizontal bars represent mean values; p-values are relative to control (0 µM) using an unpaired two-tailed t-test. **(C)** Proposed mechanism of GPR30 signaling in uterine leiomyoma: Estrogen or selective agonist G-1 activates membrane GPR30/GPER, stimulating phospholipase C (PLC) and inositol 1,4,5-trisphosphate (IP_3_) production to trigger IP_3_R-mediated calcium release from endoplasmic reticulum (ER) stores, driving cell proliferation. Pharmacological inhibition with G-15 blocks GPR30 activation, calcium mobilization, and leiomyoma growth.

### Calcium Release In HuLM Cells Is Due to GPR30

To confirm the role of GPR30 in mediating estrogen-induced calcium signaling, we utilized a selective agonist (G-1) and selective antagonist (G-15) for GPR30 (15, 16). Administration of 1 μM G-1 led to robust calcium release in 22.5 percent of cells (**Figure 4A**). Administration of 10 μM G-1 did not lead to a significantly higher increase in responders, indicating that 1 μM is saturating. This is consistent with the reported affinity of G-1 for GPR30 in the nanomolar concentration range (15). To examine whether the GPR30 antagonist G-15 can block the ability of E2 to cause calcium release, HuLM cells were pre-treated with G-15 prior to stimulation with E2. Pre-incubation with G-15 markedly attenuated calcium release by E2 (**Figure 4A; Supplementary Movie 2**). As an additional control, we examined calcium release in cells maintained in calcium-free imaging solution. Under calcium-starved conditions, cells still exhibited calcium release in response to E2 and G-1, confirming that the observed calcium signals originated from intracellular stores rather than calcium influx from the extracellular medium. This finding is consistent with GPR30 coupling to intracellular calcium release mechanisms, likely through activation of phospholipase C and generation of inositol 1,4,5-triphosphate (IP_3_), which triggers calcium release from the endoplasmic reticulum. These results indicate that GPR30 is the primary estrogen receptor responsible for rapid calcium mobilization in leiomyoma cells.

### GPR30 Antagonist G-15 Inhibits HuLM Cell Proliferation

Having established that GPR30 mediates estrogen-induced calcium release in leiomyoma cells, we next investigated whether GPR30 signaling contributes to leiomyoma cell proliferation. HuLM cells were treated with increasing concentrations of the GPR30 antagonist G-15 (0, 0.1,1, 2, 5 and 10 µM) for 48 hours, and cell viability was assessed using the MTT assay (**Figure 4B**). G-15 treatment resulted in a dose-dependent reduction in HuLM cell viability. At concentrations of 5 μM and 10 µM G-15, cell proliferation was significantly inhibited, with viability reduced by approximately 40-50% compared to vehicle-treated control cells. These findings demonstrate that pharmacological blockade of GPR30 significantly impairs leiomyoma cell proliferation, establishing a functional link between GPR30-mediated calcium signaling and leiomyoma growth (**Figure 4C**).

## DISCUSSION

This study identifies a novel non-genomic estrogen signaling pathway in human leiomyoma cells, demonstrating that estrogen drives leiomyoma cell proliferation via GPR30-mediated calcium mobilization. We show that GPR30 is abundantly expressed in HuLM cells with a predominantly intracellular localization (**Figure 1**), positioning it to mediate rapid, non-genomic estrogen responses. 17β-estradiol (E2) induces robust, dose-dependent cytosolic calcium release from intracellular stores (**Figures 2** and **3**). Selective activation of GPR30 with G-1 fully recapitulates these calcium dynamics, while GPR30 blockade with G-15 attenuates E2-stimulated calcium release and reduces HuLM cell proliferation by up to 50% (**Figure 4**). Together, these findings establish GPR30 as a primary membrane receptor coupling estrogen signaling to intracellular calcium release and leiomyoma growth. This mechanism represents a distinct pathway from classical nuclear estrogen receptor signaling and may offer new therapeutic opportunities for targeting estrogen-dependent leiomyoma growth.

These results expand growing evidence that GPR30 contributes to leiomyoma pathophysiology and environmental endocrine disruption. Previous studies identified GPR30 overexpression in leiomyoma cells (9) and implicated GPR30 in activating MAPK and EGFR pathways in fibroids (5-8). GPR30 gene polymorphisms have been associated with increased leiomyoma risk, providing further evidence for the importance of this receptor in leiomyoma development (17). In addition, calcium channel subtypes TRPC1 and TRPM7 had higher expression in uterine fibroids and surrounding smooth muscles; modification of their expression significantly inhibited cell proliferation {Kim, 2005 #2}(10). In a prior study, we demonstrated that modulating calcium signaling with simvastatin triggers robust cytosolic and mitochondrial calcium overload, leading to caspase-3 activation and apoptosis (11). Chelation of intracellular calcium with BAPTA completely abolished simvastatin-induced apoptosis, and pharmacologic blockade of L-type voltage-gated calcium channels prevented cell death (11). Our most recent findings expand on previous studies, linking GPR30 activation by estrogen to intracellular calcium mobilization, a critical second messenger that regulates cell proliferation and survival. This connection between estrogen, GPR30, and calcium provides a mechanistic framework for understanding how hormonal signals are transduced into proliferative responses in leiomyoma cells.

Targeting GPR30 presents distinct therapeutic advantages over current medical management. Standard therapies such as GnRH agonists, aromatase inhibitors, or selective estrogen receptor modulators rely on systemic hormone suppression, causing severe side effects including low bone density, vasomotor symptoms, and cardiovascular risks (1, 3). In contrast, GPR30 antagonism selectively targets rapid, membrane-initiated calcium signaling without suppressing systemic estrogen or altering classical nuclear receptor transactivation. Furthermore, because G-15 directly impairs leiomyoma cell proliferation even after cell transformation (**Figure 4B**), GPR30 inhibitors could be used therapeutically to halt active tumor growth. This finding is consistent with preclinical models demonstrating GPER-selective antagonists, such as G36, inhibits estrogen-dependent gynecologic tumor growth in xenograft models (18).

Our data indicate that GPR30 couples to intracellular calcium release through activation of phospholipase C and generation of IP_3_, which triggers calcium release from the endoplasmic reticulum (Figure 4C). This is supported by our observation that E2- and G-1-induced calcium release persists in calcium-free extracellular solution (Figure 4A), ruling out calcium influx as the primary source of the signal. Superresolution imaging confirming the intracellular localization of GPR30 (Figure 1) aligns with this model, as GPR30 on ER membranes would be ideally positioned to couple to IP_3_ receptors and modulate calcium release.

Future research should focus on identifying the immediate downstream effectors of GPR30-mediated calcium release, such as calcium-dependent kinases (CaMKII, PKC) and transcription factors (NFAT), which regulate cell cycle progression and pro-proliferative gene expression. The effect of GPR30-mediated calcium activation on downstream effects EGFR and MAPK signaling warrants further investigation (5-7). Furthermore, an essential open question is whether GPR30 antagonism alters extracellular matrix (ECM) production, a clinical hallmark of fibroids responsible for tumor bulk and symptoms (1). Because calcium signaling regulates ECM synthesis in other tissues, evaluating whether GPR30 antagonists reduces collagen and fibronectin deposition in 3D culture systems and animal models will be critical for translating these findings into clinical therapies for leiomyomas.

## ACKNOWLEDGEMENTS

Research reported in this publication was supported by the National Institute of General Medical Sciences of the National Institutes of Health under Award Number T34GM136492 (AM) and R01GM081685 (DB). The content is solely the responsibility of the authors and does not necessarily represent the official views of the National Institutes of Health.

## REFERENCES

1. Borahay MA, Al-Hendy A, Kilic GS, Boehning D. Signaling Pathways in Leiomyoma: Understanding Pathobiology and Implications for Therapy. Mol Med 2015;21:242–56.

2. De La Cruz MS, Buchanan EM. Uterine Fibroids: Diagnosis and Treatment. Am Fam Physician 2017;95:100–7.

3. Borahay MA, Asoglu MR, Mas A, Adam S, Kilic GS, Al-Hendy A. Estrogen Receptors and Signaling in Fibroids: Role in Pathobiology and Therapeutic Implications. Reprod Sci 2017;24:1235–44.

4. Prossnitz ER, Barton M. The G-protein-coupled estrogen receptor GPER in health and disease. Nat Rev Endocrinol 2011;7:715–26.

5. Jiang X, Ye X, Ma J, Li W, Wu R, Jun L. G protein-coupled estrogen receptor 1 (GPER 1) mediates estrogen-induced, proliferation of leiomyoma cells. Gynecol Endocrinol 2015;31:894–8.

6. Liu J, Yu L, Castro L, Yan Y, Sifre MI, Bortner CD et al. A nongenomic mechanism for “metalloestrogenic” effects of cadmium in human uterine leiomyoma cells through G proteincoupled estrogen receptor. Arch Toxicol 2019;93:2773–85.

7. Li Z, Lu Q, Ding B, Xu J, Shen Y. Bisphenol A promotes the proliferation of leiomyoma cells by GPR30-EGFR signaling pathway. J Obstet Gynaecol Res 2019;45:1277–85.

8. Leiber D, Burlina F, Byrne C, Robin P, Piesse C, Gonzalez L et al. The sequence Pro295-Thr311 of the hinge region of oestrogen receptor alpha is involved in ERK1/2 activation via GPR30 in leiomyoma cells. Biochem J 2015;472:97–109.

9. Tian R, Wang Z, Shi Z, Li D, Wang Y, Zhu Y et al. Differential expression of G-proteincoupled estrogen receptor-30 in human myometrial and uterine leiomyoma smooth muscle. Fertil Steril 2013;99:256–63 e3.

10. Kim BY, Cho CH, Song DK, Mun KC, Suh SI, Kim SP et al. Ciglitizone inhibits cell proliferation in human uterine leiomyoma via activation of store-operated Ca2+ channels. Am J Physiol Cell Physiol 2005;288:C389–95.

11. Borahay MA, Kilic GS, Yallampalli C, Snyder RR, Hankins GD, Al-Hendy A et al. Simvastatin potently induces calcium-dependent apoptosis of human leiomyoma cells. J Biol Chem 2014;289:35075–86.

12. Ke X, Cheng Z, Qu X, Dai H, Zhang W, Chen ZJ. High expression of calcium channel subtypes in uterine fibroid of patients. Int J Clin Exp Med 2014;7:1324–30.

13. Carney SA, Tahara H, Swartz CD, Risinger JI, He H, Moore AB et al. Immortalization of human uterine leiomyoma and myometrial cell lines after induction of telomerase activity: molecular and phenotypic characteristics. Lab Invest 2002;82:719–28.

14. Revankar CM, Cimino DF, Sklar LA, Arterburn JB, Prossnitz ER. A transmembrane intracellular estrogen receptor mediates rapid cell signaling. Science 2005;307:1625–30.

15. Bologa CG, Revankar CM, Young SM, Edwards BS, Arterburn JB, Kiselyov AS et al. Virtual and biomolecular screening converge on a selective agonist for GPR30. Nat Chem Biol 2006;2:207–12.

16. Dennis MK, Burai R, Ramesh C, Petrie WK, Alcon SN, Nayak TK et al. In vivo effects of a GPR30 antagonist. Nat Chem Biol 2009;5:421–7.

17. Kasap B, Ozturk Turhan N, Edgunlu T, Duran M, Akbaba E, Oner G. G-protein-coupled estrogen receptor-30 gene polymorphisms are associated with uterine leiomyoma risk. Bosn J Basic Med Sci 2016;16:39–45.

18. Petrie WK, Dennis MK, Hu C, Dai D, Arterburn JB, Smith HO et al. G protein-coupled estrogen receptor-selective ligands modulate endometrial tumor growth. Obstet Gynecol Int 2013;2013:472720.

